# Dominant negative FADD/MORT1 inhibits the development of intestinal intraepithelial lymphocytes with a marked defect on CD8αα+TCRγδ+ T cells

**DOI:** 10.1101/313742

**Authors:** Xuerui Zhang, Lina Huo, Lulu Song, Zhaoqing Hu, Xinran Wang, Yuheng Han, Ying Wang, Peipei Xu, Jing Zhang, Zi-Chun Hua

**Author notes:** Correspondence to: Zi-Chun Hua, and Jing Zhang. Xuerui Zhang and Lina Huo contributed equally to this work.

## Abstract

Intestinal intraepithelial lymphocytes are considered to be distinct from thymus-derived cells and are thought to derive locally from cryptopatch (CP) precursors. Although the development and homing of IELs have been studied in some details, the factors controlling their homeostasis are incompletely understood. Here, we demonstrate that FADD, a classic adaptor protein required for death-receptor-induced apoptosis, is a critical regulator of the intestinal IEL development. The mice with a dominant negative mutant of FADD (FADD-DN) display a defective localized intestinal IELs with a marked defect on CD8αα^+^TCRγδ^+^ T cells. Since Lin^-^ LPLs have been identified as precursors CP cells for CD8αα^+^ development, we analyzed lamina propria lymphocytes (LPLs) and found the massive accumulation of IL-7R^-^lin^-^ LPLs in FADD-DN mice. IL-7 plays a differentiation inducing role in the development of intestinal IELs and its receptor IL-7R is a transcriptional target of Notch1. The level of Notch1 expression also showed very low in Lin^-^ LPLs cells from FADD-DN mice compared with normal mice, indicating a possible molecular mechanism of FADD in the early IEL development. In addition, loss of γδ T-IELs induced by FADD-DN results in a worsening inflammation in murine DSS-induced colitis model, suggesting a protective role of FADD in the intestinal homeostasis.

## Introduction

Intraepithelial lymphocytes (IELs), which are integral to the intestinal mucosal associated lymphoid system, play a key role in maintaining immune homeostasis of intestine. They constitute a constellation of barrier immune cells and contribute to the intestinal function by developing tolerance to food and microbial antigens in normal physiological state and controlling insults from pathogens and deleterious tissue inflammation during mucosal infections [1, 2]. Studies conducted to date have revealed that these T cells consist of two main subpopulations. One bears either CD4 or CD8α/β molecules and TCR-α/β as thymus dependent. The other bears CD8α/α molecules and either TCR-γ/δ or TCR-α/β present in the absence of a thymus. These thymus-independent IELs like CD8α/α T cells are located mainly within the gut epithelium and develop locally [3, 4] The local development may favor the production of TCR-γ/δ cells and have a reduced capacity to induce TCR-α/β rearrangements or pre-Tα chain expression. The pre-Tα chain is an essential component of the pre-TCR that is responsible for the predominance of TCR-α/β production by the thymus. Gut intraepithelial CD8 T cells are thought to derive locally from cryptopatch (CP) precursors. The intermediate stages of differentiation between CP and mature T-IEL were not clear. There are striking differences in T cell differentiation process in the gut, when compared with T cell differentiation in the thymus, but far less is known about the molecules and signaling pathways that regulate the differentiation.

Fas-associated protein with death domain (FADD) is an adaptor protein critical for the death receptors (DRs) apoptotic signaling [5, 6]. When extrinsic apoptosis triggered, FADD interacts with death receptor (DR) like Fas or TRAIL, leading to the recruitment of procaspase-8 and its activation, then consequent apoptosis of the cells [7–11]. Besides being a main death adaptor molecule, FADD participates in other biological processes, such as embryogenesis [12], innate immunity [13], T cell activation and proliferation [14]. FADD deficiency in the immune system leads to inhibition of thymocyte development in the CD4^-^CD8^-^ (DN) stage [15]. In transgenic mice expressing a dominant negative FADD mutant (FADD-DN) under control of the lck promoter, there is also a block at this stage and a defect in the progression of thymocytes from CD25^+^CD44^-^ to CD25^-^CD44^-^ phenotype, which is associated with pre-TCR expression. Several studies on FADD-DN expression have demonstrated its suppression role on T-cell proliferation and supported an acknowledgement of the functional FADD signaling essential for normal T cell development and T cell activation. But all these studies only focused on TCR-α/β T cells from thymus dependent, this is a meaningful question whether FADD also affects the development of TCR-γ/δ T cell as thymus-independent, such as IELs.

Here we identify a novel role for FADD in the intestinal immune system. In FADD-DN transgenic mice, a significant decreased subset of CD8^+^TCRγδ^+^T cell is observed in intestinal IELs. We provide evidences that FADD-DN expression inhibits the development of CD8^+^TCRγδ^+^ T through impeding the IL-7R expression in their precursors. Loss of CD8^+^TCRγδ^+^ T as an impaired intestinal immunologic barrier, so FADD-DN mice develop more severe pathological phenotypes in DSS-induced colitis.

## Results

### Lack of γδ T cells in FADD-DN transgenic mice

To test the role of FADD in the development of murine intestinal IEL, the transgenic mice expressing a dominant negative mutant of FADD/MORT1 (FADD-DN) under control of the mouse lck proximal promoter was used in the study. FADD-DN lacks the death effector domain which is required for recruiting and activating caspase-8 during apoptosis (Fig. 1A). 16 kDa FADD-DN protein was detected by western blotting in the T-cell specific expression (Fig. 1B).

**Figure 1.**
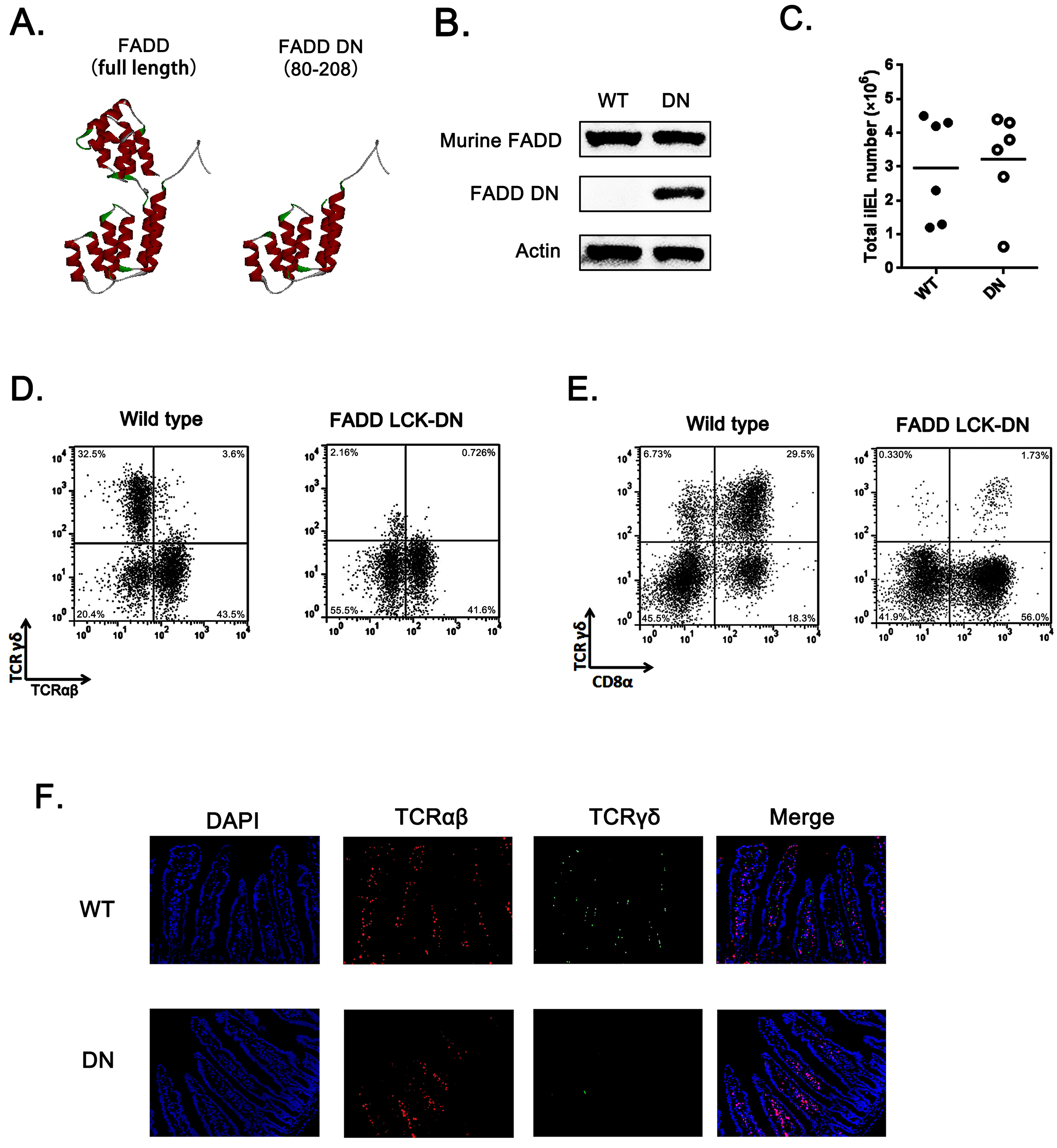
Loss of γδ T cells in FADD-DN transgenic mice. A. The three-dimensional structures of FADD [Protein Data Bank (PDB); accession number: 2GF5] and FADD-DN (death domain of FADD). B. Western blot analysis of FADD expression in wildtype (WT) and FADD-DN mice. C. Total IEL numbers of wild type and FADD-DN mice. D. FACS analysis ofγδ IELs recovered from intestines of wild type and FADD-DN mice. E. Immunofluorescent staining of γδ and αβ IELs in intestinals from wild type and FADD-DN mice.

Murine intestinal T-IELs develop by thymic and extrathymic pathways. About half derive from the peripheral lymphoid organs to the gut (TCRαβ^+^), and the other half differs from peripheral T cells, most expressing TCRγδ^+^ [16]. The local development may favor the production of TCRγδ^+^ cells. Analysis of IELs in total number showed no obvious differences in matched adult control and FADD-DN mice (Fig. 1C). When the IEL subsets were examined, TCRγδ^+^ cells was missing in FADD-DN mice (Fig. 1D and 1E). By immunofluorescence in histological sections of the small intestines, the decreased γδ T-IELs were more easier and direct to be observed. (Fig. 1F).

### A selective deficiency of CD8αα γδ T cells caused by FADD-DN expression

T-IELs consist mainly of two populations of CD8^+^ T cells. One bears CD8αβ^+^TCRαβ^+^; the other bears homodimeric α/α CD8 chains, and is mostly TCRγδ^+^ or TCRαβ^+^, it is thymo-independent [17]. Two subgroups of CD8αα^+^ and CD8αβ^+^ were isolated for further testing the proportion changes of TCRγδ^+^ or TCRαβ^+^ (Fig. 2A). In CD8αα^+^ subset, over 40% of CD8αα^+^ T expressing TCRγδ were observed in normal mice, but few γδ T cells in FADD-DN mice (Fig. 2B). No significant difference in the subset of CD8αβ^+^ αβ T was shown in both mice (Fig. 2C). By comparison of quantitative statistics, the effect of FADD-DN is mainly on the defect of CD8αα^+^ γδ T (Fig. 2D). The properties of the IEL compartment are known to change with age. Analysis of a group of aged (3 to 8 weeks old) FADD-DN and littermate control mice showed that CD8αα^+^ γδ T cells always maintained low number along the time-span in FADD-DN mice, while two groups had a similar kinetic on CD8αα^+^TCRαβ^+^ cells (Fig. 2E and 2F). These results suggest that FADD-DN expression inhibits the development of CD8αα^+^ γδ T cells.

**Figure 2.**
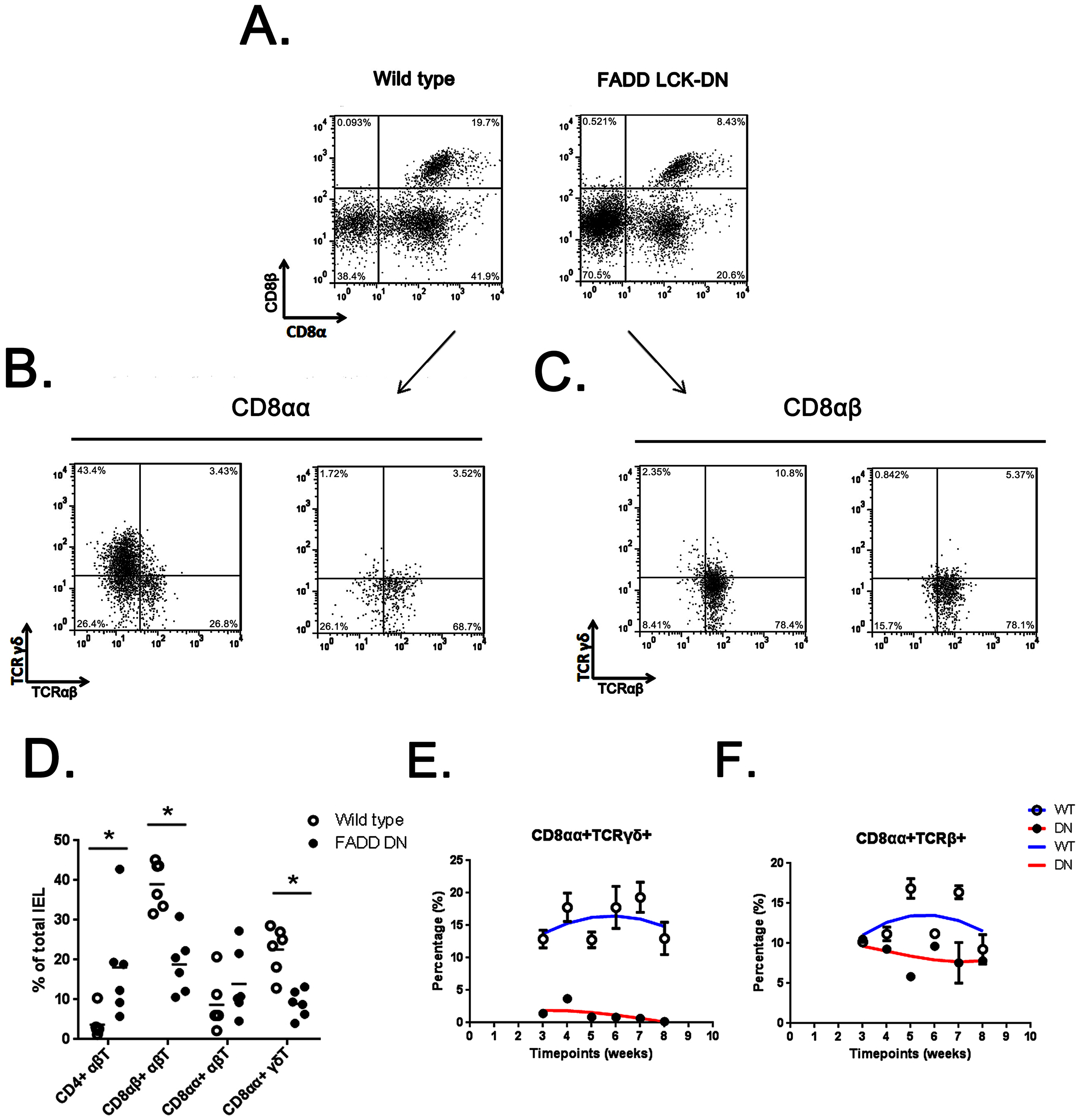
A selective deficiency of CD8αα^+^ γδ T cells in FADD-DN mice. A. FACS analysis of CD8αα^+^ and CD8αβ^+^ subsets in the IELs from wild type and FADD-DN mice. B. FACS analysis of TCRαβ^+^ and TCRγδ^+^ subsets in the CD8αα^+^ cells of IELs from wild type and FADD-DN mice. Left: wild type; Right: FADD-DN. C. FACS analysis of TCRαβ^+^ and TCRγδ^+^ subsets in the CD8αβ^+^ cells of IELs from wild type and FADD-DN mice. Left: wild type; Right: FADD-DN. D. Percentages of indicatedsubsets among total IELs from wild type and FADD-DN mice. E. Time-dependent changes of CD8αα^+^TCRγδ^+^subset in the IELs from wild type and FADD-DN mice analyzed by FACS. F. Time-dependent changes of CD8αα^+^TCRβ^+^subset in the IELs from wild type and FADD-DN mice analyzed by FACS. * P < 0.05.

### The effect of FADD-DN on intestinal IEL development is thymo-independent

T-IEL may originate from both thymus and extrathymus [4]. To distinguish the pathway for FADD-DN to regulate the development of γδ T cells, we firstly examined the percentage of CD4^-^CD8^-^γδ^+^ T cells in the thymus by FACS. In FADD-DN mice, CD4^-^CD8^-^ γδ T cells from thymus show normal compared to the wild type mice (Fig. 3A, 3B and 3C), indicating an intact development of γδ T cells in the thymus.

**Figure 3.**
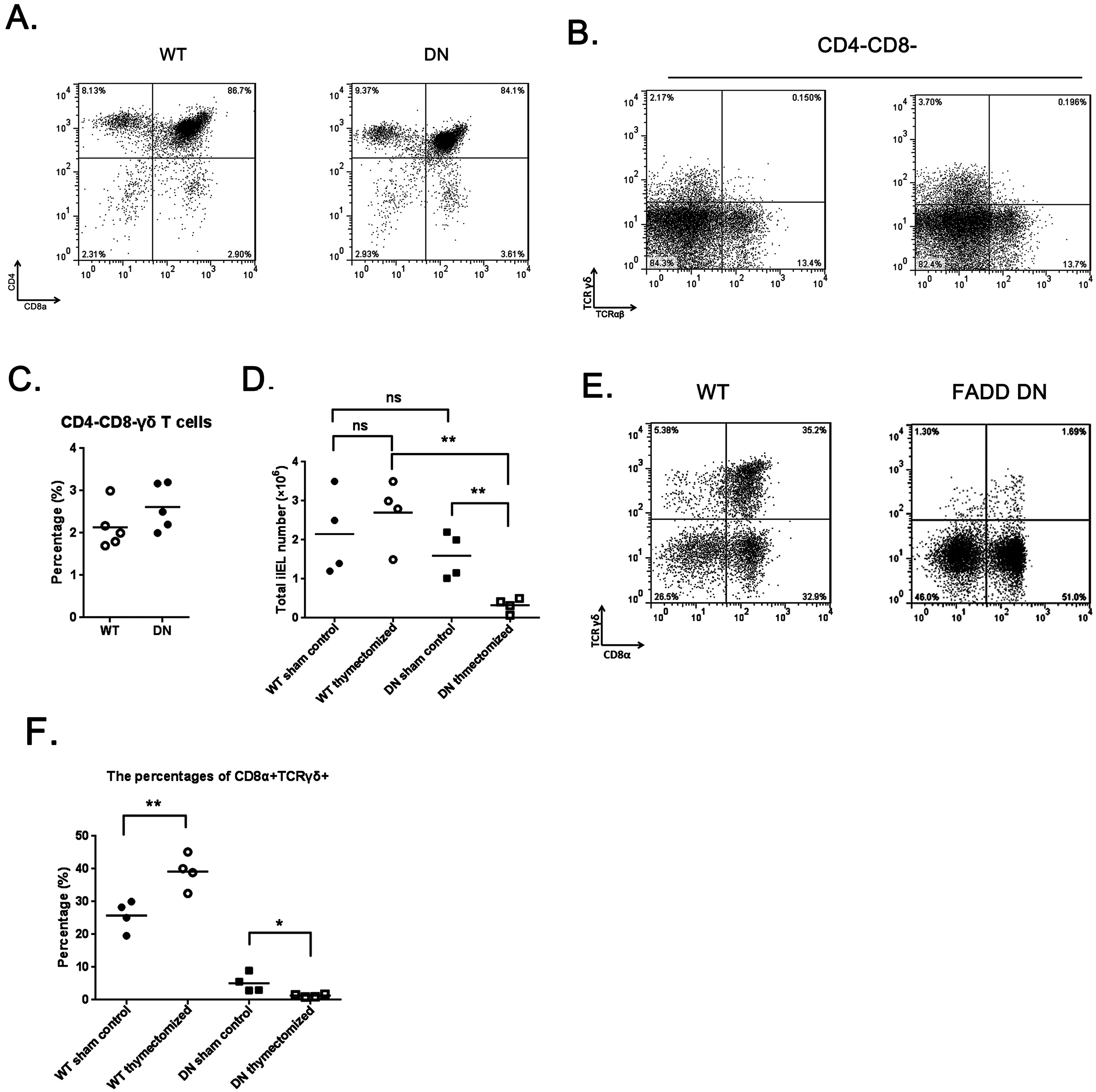
The effect of FADD-DN on intestinal IEL development. A. FACS analysis of CD4^+^ and CD8^+^ subsets in thymus from wild type and FADD-DN mice. B. FACS analysis of CD4^-^CD8^-^ γδ T cells in thymus from wild type and FADD-DN mice. C. Percentages of CD4^-^CD8^-^ γδ T cells in thymus from wild type and FADD-DN mice. D. The changes of total IEL numbers from thymecomized wild type and FADD-DN mice. E. FACS analysis of CD8^+^ γδ T cells in IELs from wild type and FADD-DN mice 4 weeks after thymectomy. F. The analysis of percentages of CD8α^+^TCRγδ^+^ subset among total IELs from thymectomized wild type and FADD-DN mice. * P < 0.05; ** P < 0.01.

In euthymic mice, competition for some local cytokines exists between αβ T cells of thymic origin and extrathymic T cell progenitors, and the extrathymic development of T-IEL is severely repressed [18]. In the athymic mice, the extrathymic lymphopoiesis emerges with a priority toward γδ T cells and with a paucity of αβ T cells [19], which will populate mainly in the intestinal mucosal leading to the accumulation of T-IELs, especially γδ T cells [20].To confirm the effect of FADD-DN on locally developmental IELs, the thymectomy was performed on both wild type and FADD-DN mice. After 4 weeks, analysis of total IELs in thymectomized mice showed a marked reduction (6-fold) in FADD-DN mice (Fig. 3D), indicating there is an impaired local development of intestinal IELs. Compared with the proportion of CD8α^+^TCRγδ^+^ T cells, thymectomized FADD-DN mice also showed missing similar to previous data (Fig. 3E). By quantifying the relative the percentage, CD8α^+^TCRγδ^+^ T showed further reduced in FADD-DN mice with thymectomy (Fig. 3F). Thus, it is reasonable to speculate that the few intestinal γδ IEL-T cells in FADD-DN mice seem to derive from thymus originally, while the local intestinal-derived γδ IEL-T cells are deficient. Taken together, the deficiency of γδ IEL-T cells caused by FADD-DN expression might be stunting locally development.

### The development of IELs is arrested at stage of Lin^-^ LPLs in FADD-DN mice

Murine intestinal IEL originate from their own pre-existing stem cells present in the intestinal (i.e. cryptopatches) [4, 21, 22] and appendix [23]. Intestinal T cell precursors have specific phenotype: lineage markers negative (Lin^-^) Thy^+^c-kit^+^IL-7R^+^CD44^+^CD25^+^ [4]. To determine at which point the differentiation of murine intestinal IELs is plagued by FADD-DN, we examined the Lin^-^ cell types prepared from IELs and lamina propria lymphocytes (LPLs).

By flow cytometry analysis with lineage markers, the number of Lin^-^ IELs from FADD-DN showed double (Fig. 4A). Gating on Lin^-^ IELs for further CD8α expression, compared with wild type mice, the IELs subgroup of CD8^+^Lin^-^ was deficient in FADD-DN mice (Fig. 4B), suggesting the differentiation of IELs was blocked in Lin^-^CD8^-^. For a better understanding of IEL maturation in earlier events, we examined Lin^-^ LPLs which have been proved equal to precursors Lin^-^ CP. Notably, a massive accumulation of Lin^-^ LPLs was observed in FADD-DN mice, an clear hint on the attested stage in the development (Fig. 4C). Our previous research found that FADD regulates thymocyte development at the β-selection checkpoint by modulating Notch signaling [7]. Notch1 is essential for T differentiation and specifying the cell fate, so the Notch1 transcription in progenitor Lin^-^ LPLs was detected. Excitedly, there are two groups in normal Lin^-^ LPLs with different forms of Notch expression: high expression and low expression, but in FADD-DN Lin^-^ LPLs, the group with high expression of Notch is obviously missing (Fig. 4D). Notch1 controls T-cell development in part by regulating the stage- and lineage-specific expression of IL-7R which is a transcriptional target of Notch1. C-Kit is a tyrosine kinase receptor and extrathymus-derived IELs normally seen in older mice were c-Kit-dependent and represented a distinct lineage of T cells with unique developmental and functional attributes. Then two crucial markers: CD117 (c-kit) and CD127 (IL-7R) were checked by FACS. By comparisons of the Lin^-^ LPL subsets divided with c-kit and IL-7R, a clear distinct difference in IL-7R expression (CD127) was observed (Fig. 4E). There is no expression of IL-7R in Lin^-^ LPL cells from FADD-DN mice, which is well consistent with the low expression of Notch1. The enterocyte-produced IL-7 plays a differentiation inducing role in the development of intestinal IELs [24, 25]. In order to fully development, the thymic-independent TCRγδ^+^ IELs in an immature state localized in the intestine must interact with their appropriate ligands in situ, so lack of IL-7R in Lin^-^ LPL cells from FADD-DN mice might provides a novel role for FADD function in early T cell development.

**Figure 4.**
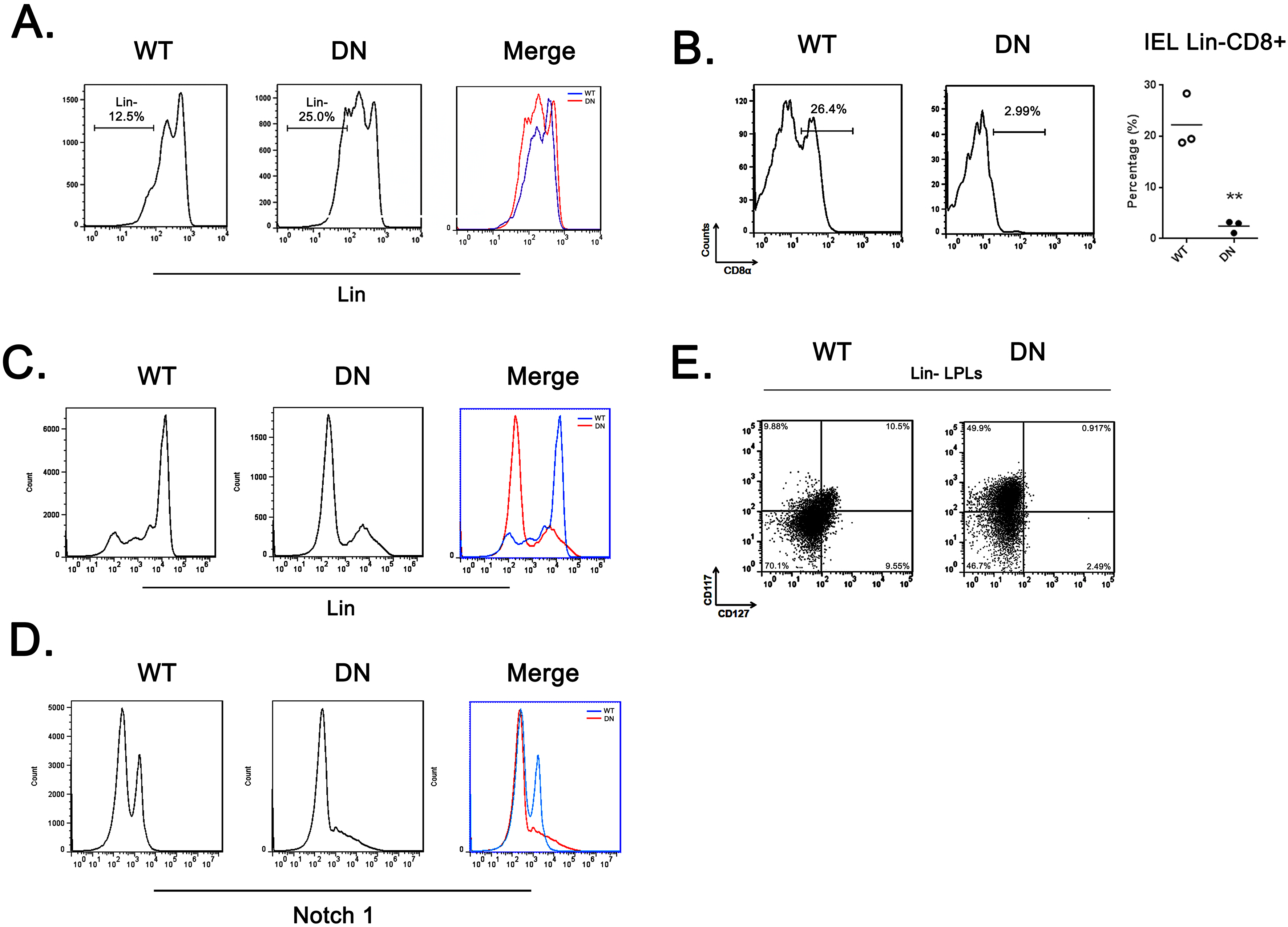
The development of IELs arrested by FADD-DN in lin^-^ LPLs with IL-7R deficiency. A. FACS analysis of lin^-^ IELs recovered from indicated groups. IELs recovered from wild type and FADD-DN mice were stained with antibodies against lineage markers (lin). B. FACS analysis of CD8α^+^ subset in the lin^-^ IELs from indicated groups. C. FACS analysis of lin^-^ LPLs recovered from indicated groups. LPLs recovered from wild type and FADD-DN mice were also stained with antibodies against lineage markers. D. FACS analysis of CD117^+^CD127^+^ subset in the lin^-^ LPLs from indicated groups. E. FACS analysis of Notch1 expression in the LPLs from indicated groups.

### The role of FADD-DN in intestinal immunoregulation

As the first-line defense against infectious agents, γδ T-IELs are involved in intestinal immunoregulation [26]. Depletion or deficiency in γδ T cells aggravates intestinal inflammation in almost every investigated model, especially DSS-induced colitis [27]. Our finding about loss of γδ T-IELs in FADD-DN mice draws substantial experimental interest on its responses in pathological condition. DSS-induced colitis model was established in both wild type and FADD-DN mice. Severe illness, which was characterized by diarrhea, intestinal bleeding, body weight loss and shortened colon length, was observed at about 4 days after DSS administration. Compared with wild type group, FADD-DN mice exhibited severe colitis with substantial reduction in colon length (Fig. 5A and 5B). From 3 days to 7 days of DSS administration, FADD-DN mice showed significant loss of body weight compared to the wild type group (Fig. 5C), and its disease activity index (DAI) also increased more dramatically (Fig. 5D). The histological results demonstrated more severe pathological changes, including loss of goblet cells, distortion of crypts, and mucosal damage and necrosis in the colon specimens in DSS-treated FADD-DN mice, with evidently higher scores than wild type mice (Fig. 5E and 5F). Consistent with these phenotypes, higher levels of TNF-α, IFN-γ and IL-6 were detected in the serum of FADD-DN mice, while no significant differences in IL-12p40 and IL-1β (Fig. 5G). It is worth mentioning that FADD-DN sham control group with no colitis, also exhibited a higher level of some cytokines than wild type sham group (TNF-α, IL-12p40, and IL-1β), sugguesting FADD-DN expression might affect the gut immunity under physiological conditions.

**Figure 5.**
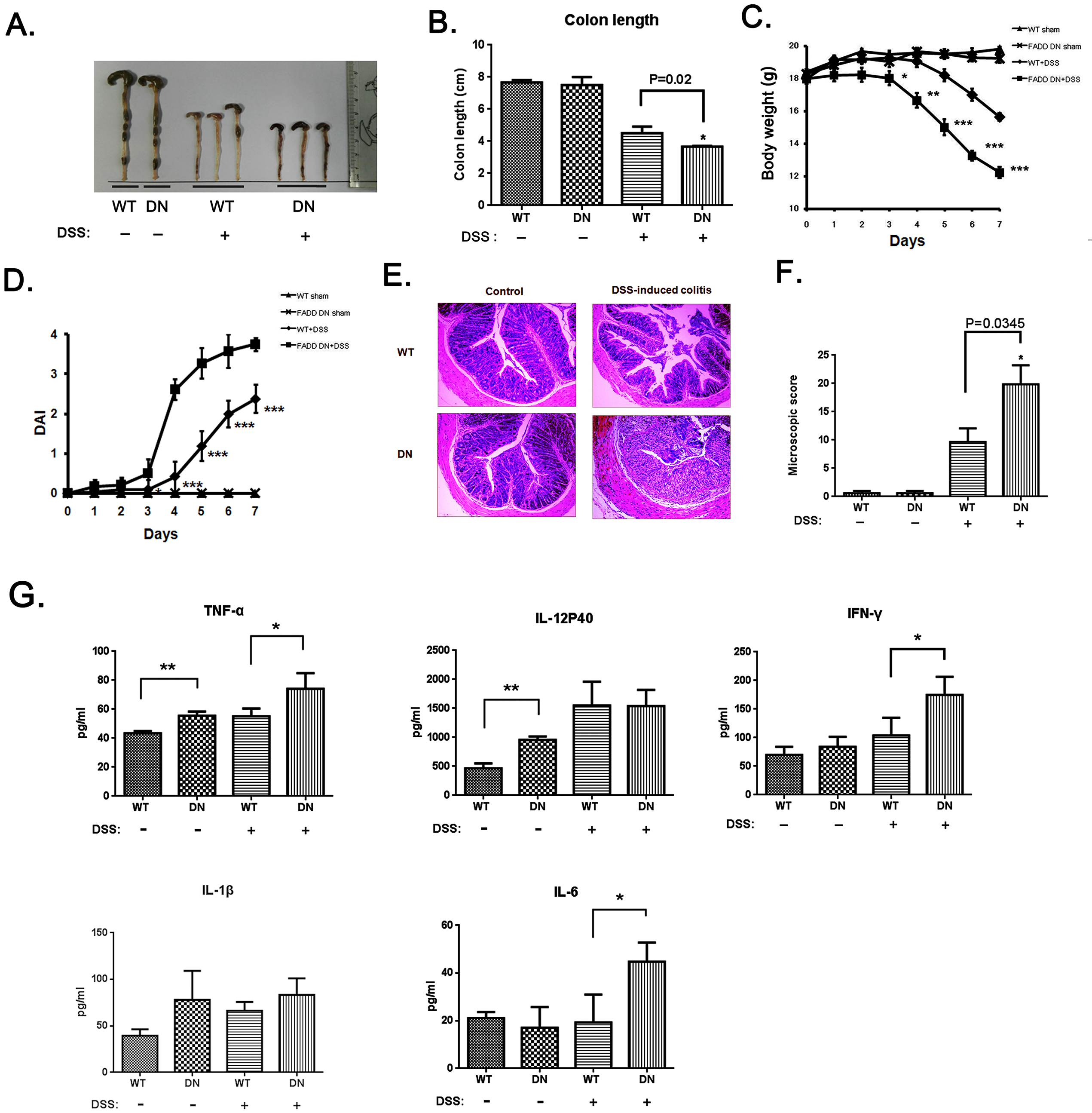
More severe DSS-induced colitis in FADD-DN mice. Mice were given 3% DSS in theirdrinking waterfor 5 days and then provided with water for another 2 days before being sacrificed. A, B. (A) Macroscopic appearances and (B) colon lengths of the mice were measured. C. Changes in the body weights of mice in each group during the experiment (n=6 per group). D. Disease activity index (DAI) was calculated (n=6 per group). E. Colon sections of indicated groups were stained with H&E. The original amplification was 100×. F. Histological scores of colon sections from different groups were calculated as described in the Materials and methods. G. Serum levels of indicated cytokines in different groups detected by ELISA. Datashown here are from a representative experiment repeated three times with similar results.*P<0.05,**P<0.01, ***P<0.001 vs DSS-treated vehicle group.

## Discussion

To date, a plethora of reports has clearly identified FADD as an essential adaptor between death receptors and caspase-8 [28, 29]. Non-apoptotic functions of FADD have been implicated in T lymphocyte development and function, mostly in thymic αβ T cells. To clarify the role of FADD in the development of TCRγδ^+^ T cells, FADD-DN mice provided a useful tool to dissect T cell development in the mucosa for thymic and extrathymic pathways. Based on our findings, FADD play an important role for IELs development, especially for γδ^+^ T-IELs which are thought to develop locally and largely via an extrathymic pathway.

T-IEL play a critical role in regulating intestinal mucosal immune responses. γδ IEL is a crucial protective T cell subset against colitis, which has been proved in Cδ^-/-^ mice [30]. Loss of γδ IEL in FADD-DN mice is a main phenotype. The decreased CD8α^+^TCRγδ^+^ T will take responsibility for the reduced homeostasis, so FADD-DN mice showed severe inflammation in DSS-induced colitis model (Fig. 5). To track the differentiation of CD8α^+^TCRγδ^+^ T cells, we prepared intestinal T cell precursors referring to Lambolez’ paper published on J. Exp. Med. The Lin^-^ LPLs cells (or namely Lin^-^ WL) isolated by their method had been fully demonstrated to be in fact cryptopatch (CP) precursors. They all have specific phenotype: (Lin^-^) Thy^+^c-kit^+^IL-7R^+^CD44^+^CD25^+^. Analysis of Lin^-^ IELs and Lin^-^ LPLs from FADD-DN mice revealed an arrest for CD8α^+^TCRγδ^+^ T development at stage of IL-7R^-^c-kit^+^lin^-^ LPLs.The IL-7 receptor is expressed on lymphoid T and B precursors, and innate lymphoid cells. At physiological levels, IL-7 is integral to T and B cell development in primary lymphoid organs and plays an essential role in supporting normal T cell development and homeostasis. The loss of IL-7R expression in gut precursors might be responsible for the defects on IEL development in FADD-DN mice because the localized development of IEL is IL-7-dependent. The possible molecular mechanism is the deficiency of IL-7R expression in c-kit^+^lin^-^ LPLs due to the Notch1 expression suppressed by FADD-DN (Fig.4).

We previously reported that FADD regulates thymocyte development at the β-selection checkpoint by modulating Notch signaling. Notch1 mRNA increases from early thymocyte progenitor to the DN3a stage and markedly decreases at the DN3b stage commensurate with pre-TCR signaling. Differential expression of Notch represents a distinct lineage of T cells with unique developmental and functional attributes. In normal mice, we also detect two states of Notch1 expression in gut T cell precursors, but this regulation is destroyed by FADD-DN expression (Fig. 4). The effect of FADD-DN on the intestinal IELs reveals the critical nature of FADD. A Notch1 transcriptional repressor NKAP was previously identified by us to associate with FADD, suggesting a potential role of FADD in maintaining the stabilization of NKAP to inhibit Notch on transcriptional level. Besides being cytoplasmic, the FADD protein is also expressed in the nucleus of many types of cells and nuclear FADD is implicated to be involved in a functional transcription factor complex to modulate Notch expression. The regulation of FADD on Notch in the intestine IELs is under investigating, which will give a comprehensive understanding of FADD function in the physiology of immune system.

In summary, the present study demonstrates an novel function of FADD in protective T cell subset against colitis. This knowledge will enable further investigation of the role of FADD in the IELs development in order to develop novel therapeutic strategies.

## Materials and methods

### Mice

Female wild-type mice and FADD DN mice (6-8weeks) were provided by the laboratory of Aster Winoto at the University of California, Berkeley and maintained in pathogen-free conditions.

### Isolation of intraepithelial lymphocytes

Islation of intestinal Intraepithelial lymphocytes(IEL)was performed as described previously [3]. After washing the small intestine to remove fecal content, Peyer’s patches and fat tissue were extirpated. The small intestine was opened longitudinally, cut into small pieces, and washed with PBS containing 0.1% BSA, 100 U/ml penicillin and 100 μg/ml streptomycin three times. The intestines were then shaken with pre-warmed DMEM containing penicillin, streptomycin, and 5% FCS for 30 min at 225 rpm, 37°C. After filtrating the supernatants through a nylon mesh, IELs were collected by a 44-67% Percoll density gradient (Solarbio, Beijing, China) and the cells that layered between the 44-67%. After washed by PBS, cells were counted and used for the following assays.

### Isolation of lamina propria lymphocytes (LPL)

After IEL isolation, tissues were digested in RPMI1640 supplemented with 0.5 mg/ml collagenase type VIII (Sigma, USA), 0.1 mg/ml DNase I (Sigma, USA), and 10% FCS at 200 rpm, 37°C for 30 min. Lamina propria lymphocytes (LPL) were released and then subjected to Percoll fractionation as described above for isolation of IEL.

### Dextran Sodium Sulfate(DSS)-induced colitis

FADD DN or wild-type mice were fed with water charged with 3.0% (W/V) DSS S (dextran sulfate sodium salt, 36-50 kDa, 0216011080, MP Biomeicals) for 5 days and then with normal drinking water for 2 days till sacrificed. Mice given normal water served as control. Body weight of mice was recorded daily. Mice were sacrificed 7 days after DSS exposure. The length of the colons for each group were measured. After washing by PBS, colons tissue were cut and fixed in 4% (v/v) formaldehyde for histological analysis.

### Evaluation of disease activity

During the experiment, body weight of mice for each group was recorded daily and feces were collected. Disease activity index (DAI) was determined by loss of weight, stool consistency and fecal blood. The calculation of DAI was performed as previously [20].

### Histological analysis

The colons were fixed in formaldehyde, embedded in paraffin and cut into serial sections of 3-μm-thick for histological analysis. Stained with haematoxylin and eosin (H&E), tissues were analyzed by histological grading according to the criteria described previously [21].

### Immunofluorescence assay

The deparaffinized colon tissues underwent antigen retrieval and were blocked with 10% goat serum. Sections were incubated with anti-mouse TCR-αβ and TCR-γδ primary antibodies (1:50, Cell Signaling Technology (CST), Danvers, MA, USA) overnight at 4°C. After washing with PBS three times for 5 minutes, sections were incubated with secondary antibodies anti-mouse HRP (1:100) and anti-rabbit HRP (1:100) at room temperature for 30 minutes. Following washed with PBS three times for 5 minutes, slides were counterstained with DAPI (invitrogen). All images of the slides were visualized and captured by fluorescence microscope.

### Western Blotting Assay

The total proteins of splenocytes were prepared by using RIPA buffer. Samples of protein lysates were separated by 10% SDS-PAGE and electro-transferred to PVDF membrane. After blocking with 5% non-fat milk for 2 h at room temperature, membranes were incubated with against mouse FADD (1 : 1000; catalogue number ab124812; Abcam, Cambridge, UK), human FADD C-terminus (1 : 1000; 610399; BD Pharmingen, Franklin Lake, NJ, USA), β-actin (1 : 1000; AM1021B; Abgent, San Diego, CA, USA) followed by suitable secondary antibody conjugated with horseradish peroxidase (1 : 2000; Jackson Immunoresearch Laboratories, West Grove, PA, USA). Reactive bands were detected with by enhanced chemiluminescence (Tanon, Shanghai, China) and visualized with an imaging program (Tanon, Shanghai, China). The intensities of the scanned bands were normalized to the β-actin signal. Anti-mouse FADD (ab124812; Abcam) was used to detect the endogenous murine FADD, while anti-human FADD C-terminus (610399; BD Pharmingen) was used to detect FADD-DN.

### Cytokine measurement

Cytokines in serum samples including TNF-α, IL-6, IL-17, IL-1β,IL-12 were measured using ELISA kits (R&D Systems, US) according to the manufacturer’s instructions.

### Flow cytometry analysis

Cells were stained using standard procedures for surface markers. The following monoclonal antibodies were used for flow cytometry: anti-CD3 (145-2C11), anti-CD19 (1D3), anti-TCRαβ (H57-597), anti-TCRγδ (GL3), anti-erythroid cells (TER-119), anti-GR1 (8C5), anti–CD11b (M170), anti-IgM (LL41), and anti-CD8b (eBioH35-17.2). All the antibodies were purchased from BD Pharmingen (San Diego, CA). CD8αα cells were gated as CD8α^+^CD8β^-^.

### Thymectomy

Thymectomy was performed on 8-wk-old mice as described before [22]. A ventilator was used to keep mice breathing during the operation. Completeness of thymectomy was confirmed by visual inspection, both directly after removal of the organ and at the end of the experiment. Only fully thymectomized animals were included in this study.

### Statistical analysis

The experimental data were presented as mean ± SEM. The statistical significance of the differences between groups was evaluated by One-way ANOVA and Student’s t test (P< 0.05).

## Acknowledgements

This study was supported in part by grants from the Chinese National Natural Sciences Foundation (81630092, 81773099, 81570790, 81573338), the National Key R&D Research Program by Ministry of Science and Technology (2017YFA0506002, 2017YFA0104301, 2016YFC0902700) and Shenzhen Science and Technology Innovation Committee (JCYJ20160331152141936), Shenzhen Peacock Plan (KQTD20140630165057031).

## Conflict of interest

The authors indicate no potential conflicts of interest.

## References

1. Cheroutre H, Lambolez F and Mucida D. The light and dark sides of intestinal intraepithelial lymphocytes. Nature reviews Immunology. 2011; 11(7):445–456.

2. Wang X, Sumida H and Cyster JG. GPR18 is required for a normal CD8alphaalpha intestinal intraepithelial lymphocyte compartment. The Journal of experimental medicine. 2014; 211(12):2351–2359.

3. Rocha B, Vassalli P and Guy-Grand D. Thymic and extrathymic origins of gut intraepithelial lymphocyte populations in mice. The Journal of experimental medicine. 1994; 180(2):681–686.

4. Lambolez F, Azogui O, Joret AM, Garcia C, von Boehmer H, Di Santo J, Ezine S and Rocha B. Characterization of T cell differentiation in the murine gut. The Journal of experimental medicine. 2002; 195(4):437–449.

5. Newton K, Harris AW, Bath ML, Smith KG and Strasser A. A dominant interfering mutant of FADD/MORT1 enhances deletion of autoreactive thymocytes and inhibits proliferation of mature T lymphocytes. The EMBO journal. 1998; 17(3):706–718.

6. Yeh WC, de la Pompa JL, McCurrach ME, Shu HB, Elia AJ, Shahinian A, Ng M, Wakeham A, Khoo W, Mitchell K, El-Deiry WS, Lowe SW, Goeddel DV and Mak TW. FADD: essential for embryo development and signaling from some, but not all, inducers of apoptosis. Science. 1998; 279(5358):1954–1958.

7. Zhang X, Dong X, Wang H, Li J, Yang B, Zhang J and Hua ZC. FADD regulates thymocyte development at the beta-selection checkpoint by modulating Notch signaling. Cell death & disease. 2014; 5:e1273.

8. Chinnaiyan AM, O’Rourke K, Tewari M and Dixit VM. FADD, a novel death domain-containing protein, interacts with the death domain of Fas and initiates apoptosis. Cell. 1995; 81(4):505–512.

9. Zhang J and Winoto A. A mouse Fas-associated protein with homology to the human Mort1/FADD protein is essential for Fas-induced apoptosis. Molecular and cellular biology. 1996; 16(6):2756–2763.

10. Chinnaiyan AM, O’Rourke K, Yu GL, Lyons RH, Garg M, Duan DR, Xing L, Gentz R, Ni J and Dixit VM. Signal transduction by DR3, a death domain-containing receptor related to TNFR-1 and CD95. Science. 1996; 274(5289):990–992.

11. Hsu H, Shu HB, Pan MG and Goeddel DV. TRADD-TRAF2 and TRADD-FADD interactions define two distinct TNF receptor 1 signal transduction pathways. Cell. 1996; 84(2):299–308.

12. Yeh W-C, de la Pompa JL, McCurrach ME, Shu H-B, Elia AJ, Shahinian A, Ng M, Wakeham A, Khoo W and Mitchell K. FADD: essential for embryo development and signaling from some, but not all, inducers of apoptosis. Science. 1998; 279(5358):1954–1958.

13. Balachandran S, Thomas E and Barber GN. A FADD-dependent innate immune mechanism in mammalian cells. Nature. 2004; 432(7015):401–405.

14. Newton K, Harris AW, Bath ML, Smith KG and Strasser A. A dominant interfering mutant of FADD/MORT1 enhances deletion of autoreactive thymocytes and inhibits proliferation of mature T lymphocytes. The EMBO Journal. 1998; 17(3):706–718.

15. Hua ZC, Sohn SJ, Kang C, Cado D and Winoto A. A function of Fas-associated death domain protein in cell cycle progression localized to a single amino acid at its C-terminal region. Immunity. 2003; 18(4):513–521.

16. Peaudecerf L, dos Santos PR, Boudil A, Ezine S, Pardigon N and Rocha B. The role of the gut as a primary lymphoid organ: CD8 alpha alpha intraepithelial T lymphocytes in euthymic mice derive from very immature CD44(+) thymocyte precursors. Mucosal Immunol. 2011; 4(1):93–101.

17. Guy-Grand D, Cerf-Bensussan N, Malissen B, Malassis-Seris M, Briottet C and Vassalli P. Two gut intraepithelial CD8+ lymphocyte populations with different T cell receptors: a role for the gut epithelium in T cell differentiation. The Journal of experimental medicine. 1991; 173(2):471–481.

18. Jameson SC. Maintaining the norm: T-cell homeostasis. Nature reviews Immunology. 2002; 2(8):547–556.

19. Delphine Guy-Grand OA, Susanna Celli, Sylvie Darche, Michel C. Nussenzweig, Philippe Kourilsky, Pierre Vassalli. Extrathymic T Cell Lymphopoiesis. JEM. 2003; 197(3):9.

20. Guy-Grand D and Vassalli P. Gut intraepithelial lymphocyte development. Current opinion in immunology. 2002; 14(2):255–259.

21. Sugahara S, Shimizu T, Yoshida Y, Aiba T, Yamagiwa S, Asakura H and Abo T. Extrathymic derivation of gut lymphocytes in parabiotic mice. Immunology. 1999; 96(1):57–65.

22. Saito H, Kanamori Y, Takemori T, Nariuchi H, Kubota E, Takahashi-Iwanaga H, Iwanaga T and Ishikawa H. Generation of intestinal T cells from progenitors residing in gut cryptopatches. Science. 1998; 280(5361):275–278.

23. Yamagiwa S, Sugahara S, Shimizu T, Iwanaga T, Yoshida Y, Honda S, Watanabe H, Suzuki K, Asakura H and Abo T. The primary site of CD4- 8- B220+ alphabeta T cells in lpr mice: the appendix in normal mice. Journal of immunology. 1998; 160(6):2665–2674.

24. Suzuki K, Oida T, Hamada H, Hitotsumatsu O, Watanabe M, Hibi T, Yamamoto H, Kubota E, Kaminogawa S and Ishikawa H. Gut cryptopatches: direct evidence of extrathymic anatomical sites for intestinal T lymphopoiesis. Immunity. 2000; 13(5):691–702.

25. Yang H, Spencer AU and Teitelbaum DH. Interleukin-7 administration alters intestinal intraepithelial lymphocyte phenotype and function in vivo. Cytokine. 2005; 31(6):419–428.

26. Smith AL and Hayday AC. An alphabeta T-cell-independent immunoprotective response towards gut coccidia is supported by gammadelta cells. Immunology. 2000; 101(3):325–332.

27. Kuhl AA, Pawlowski NN, Grollich K, Loddenkemper C, Zeitz M and Hoffmann JC. Aggravation of intestinal inflammation by depletion/deficiency of gammadelta T cells in different types of IBD animal models. Journal of leukocyte biology. 2007; 81(1):168–175.

28. Nogusa S, Thapa RJ, Dillon CP, Liedmann S, Oguin TH, 3rd, Ingram JP, Rodriguez DA, Kosoff R, Sharma S, Sturm O, Verbist K, Gough PJ, Bertin J, Hartmann BM, Sealfon SC, Kaiser WJ, et al. RIPK3 Activates Parallel Pathways of MLKL-Driven Necroptosis and FADD-Mediated Apoptosis to Protect against Influenza A Virus. Cell host & microbe. 2016; 20(1):13–24.

29. Wang PX, Ji YX, Zhang XJ, Zhao LP, Yan ZZ, Zhang P, Shen LJ, Yang X, Fang J, Tian S, Zhu XY, Gong J, Zhang X, Wei QF, Wang Y, Li J, et al. Targeting CASP8 and FADD-like apoptosis regulator ameliorates nonalcoholic steatohepatitis in mice and nonhuman primates. Nature medicine. 2017; 23(4):439–449.

30. Inagaki-Ohara K, Chinen T, Matsuzaki G, Sasaki A, Sakamoto Y, Hiromatsu K, Nakamura-Uchiyama F, Nawa Y and Yoshimura A. Mucosal T cells bearing TCRgammadelta play a protective role in intestinal inflammation. Journal of immunology. 2004; 173(2):1390–1398.

